# Age Prediction based on Blood DNA Methylation

**DOI:** 10.1101/2024.10.25.620181

**Authors:** Liu Rui, Fan Pu, Zhang Jing, Liu Zhao, Mi Hao-yuan

## Abstract

**Objective:** In the judicial field, traditional DNA methylation age prediction models have low accuracy and poor stability. Additionally, the use of linear regression models for detection is inefficient and costly. This study aims to utilize the prediction principles of the Support Vector Regression (SVR) model, based on preliminary laboratory data from blood DNA methylation detection using the Illumina 850K chip. By selecting low-dimensional and highly linear loci, we aim to establish a highly stable and accurate blood DNA methylation age prediction model.

**Methods:** This research is based on Illumina 850K chip technology. We conducted a literature review to select CpG sites and related primers, then employed SVR for model construction and age prediction. The model was built on the Matlab2022a platform. Standard parameters were selected, and optimal values for C and g were determined using grid search and cross-validation methods. During data processing, numerical values were normalized before calculation and de-normalized to obtain the predicted values.

**Results:** The constructed model achieved an R^2^ of 0.91563 and a Mean Absolute Error (MAE) of 2.77 years. This indicates that the prediction accuracy for blood samples reached 91.56%, with an error of 2.77 years. Moreover, the accuracy of the model’s predictions decreases with increasing age.

## 1. Introduction

DNA methylation age estimation is based on the dynamic changes of DNA methylation patterns throughout the lifespan. By comparing an individual’s actual age with the level of DNA methylation in their genome, predictive models can be established to accurately infer age. DNA methylation age estimation holds broad applications and scientific significance. It can assess physiological and health conditions, explore associations between DNA methylation and aging, disease risks, and provide references for personalized medicine and health management. For instance, by comparing DNAm age with chronological age, potential health risks can be identified early, enabling appropriate preventive measures. Analyzing differences in DNAm age among different groups can reveal the impact of environmental factors (such as lifestyle, diet, and pollution) on biological age, thus informing health policies and interventions.

In forensic practice, DNA methylation age estimation offers a novel approach for forensic medicine and individual identification. Analyzing DNA methylation levels to deduce biological age provides crucial information in legal contexts. Firstly, DNAm age can aid in criminal investigations by accurately determining the age of suspects using DNA methylation analysis of extracted samples. Secondly, it facilitates age determination in cases involving unidentified individuals, such as forensic examinations of bodies or child abduction cases. Thirdly, DNAm age can assist in defining criminal responsibility age, which varies between minors and adults in many legal systems.

While DNA methylation holds significant potential in forensic applications, rigorous research and practical experience are essential to ensure its scientific validity, stability, and accuracy in serving judicial practices objectively and fairly.

With the advancing field of epigenetics, the focus in human characterization studies is shifting towards single nucleotide polymorphisms (SNPs), alongside traditional markers like restriction fragment length polymorphisms (RFLP) and short tandem repeats (STR). DNA methylation, as a crucial epigenetic marker, complements these markers and plays a pivotal role in age estimation models based on its dynamic changes across the lifespan. Traditional methylation models typically employ linear regression requiring multiple markers and samples, thereby increasing complexity and costs. This study explores SVR modeling principles to predict age, utilizing fewer but stronger linear markers, aiming to establish a highly stable and accurate blood DNA methylation age estimation model.

Blood-based DNA methylation age estimation has been a prominent research area. Previous studies have utilized technologies like Illumina Infinium 450K chips and EpiTYPER systems to establish age prediction models based on various CpG sites, demonstrating different levels of mean absolute deviation (MAD). The evolution of these models continues with improvements in SVR models, aiming for robustness and compatibility in age prediction from biological samples.

In this study, a comparative analysis between multiple linear regression (MLR) and SVR models using sample data suggests that SVR models provide higher repeatability, better fit, and greater compatibility for developing accurate age prediction models based on blood DNA. Through literature review and analysis, this research aims to select CpG sites common in saliva and blood DNA, analyze their methylation levels using Illumina 850K chip technology, and establish a SVR regression model for blood-based DNA methylation age estimation.

## 2. Methods

### 2.1 Sample Collection

This study utilized Illumina 850K chip data from 80 unrelated individual blood samples collected and tested by the Ministry of Public Security Appraisal Center. All samples were collected with informed consent from the individuals and approved by the ethics committee of the Ministry of Public Security Appraisal Center. The age of blood samples ranged from 20 to 59 years, with an average age of 39.08 years, as shown in Tab 1.

**Tab.1.** Blood sample distribution information.

| Age groups | Sample size | Average age |
| --- | --- | --- |
| 20 ~ 29 | 20 | 23.55 |
| 30 ~ 39 | 20 | 34.75 |
| 40 ~ 49 | 20 | 44.45 |
| 50 ~ 59 | 20 | 53.55 |

### 2.2 Selection of CpG Sites

Based on literature review and laboratory data from the Ministry of Public Security Appraisal Center, 5 CpG sites potentially associated with age were initially selected as candidate sites for model training and research. These sites were numbered accordingly (as shown in Tab 2).

**Tab.2.** Initial screening site number information.

| Identification | SITE |
| --- | --- |
| CpG1 | CG21572722 |
| CpG2 | CG19283806 |
| CpG3 | CG03607117 |
| CpG4 | CG13552692 |
| CpG5 | CG14692377 |
| CpG6 | CG25410668 |
| CpG7 | CG04875128 |
| CpG8 | CG13552692 |

### 2.3 Pearson Correlation Analysis

Pearson correlation analysis is a statistical method to assess the linear relationship and direction between two variables. In this experiment, the “cor()” function in RStudio was used to analyze data from 80 samples across 5 candidate CpG sites detected by the 850K chip. This analysis evaluated the correlation between methylation levels at each candidate site and the corresponding age of the samples, yielding correlation coefficients (denoted as “r”) and regression scatter plots.

#### 2.4 Selection and Optimization of Site Data

To meet the significant correlation requirement between variables using Support Vector Machine (SVM), CpG sites with |r| > 0.5 from Pearson correlation analysis were selected for subsequent SVR regression model training and construction. Outlier samples were identified and excluded during data optimization to enhance prediction accuracy of the model. Normalization of selected sample data based on MATLAB R2022a’s “mapminmax(‘apply’)” function was performed to mitigate the impact of differing magnitude levels on model performance and computation speed. The final prediction values were reverse-normalized using “mapminmax(‘reverse’)” to obtain age predictions in standard numerical ranges.

### 2.5 Construction of Age prediction Model

Samples, after selection, were randomly shuffled with 80% used for training and 20% for testing the model. The SVR regression model was built using MATLAB R2022a and the libsvm-3.34 toolkit, employing grid search and cross-validation to optimize parameters C (regularization parameter) and γ (gamma). The “svmtrain()” function established the model for predicting age from methylation levels at CpG sites, while “svmpredict()” simulated age predictions based on methylation levels in the test set. Visualization and mathematical analysis of model performance were conducted through plotting and calculation of relevant error metrics.

### 2.6 Analysis of Model Accuracy

The accuracy of the age inference model was analyzed across four age groups (20-29, 30-39, 40-49, 50-59 years) using 76 selected blood samples. Model performance indicators such as Mean Absolute Deviation (MAD) and accuracy were computed, with an error margin set at ±5 years considered acceptable.

## 3. Results

### 3.1 Pearson Correlation Analysis

Based on Illumina 850K chip data from 80 blood samples, DNA methylation levels at 8 CpG sites were correlated with sample age, as depicted in scatter plots in Fig. 1. The calculated Pearson correlation coefficients are presented in Tab. 3. Among the 8 candidate sites, CpG2, CpG3, CpG4, CpG5, and CpG8 showed |r| values above 0.5, indicating a strong positive correlation between methylation levels and sample age. CpG8, while more correlated than CpG5, exhibited more outliers, which were unfavorable for SVR model construction. Therefore, CpG2, CpG3, CpG4, and CpG5 were chosen for the SVR model.

**Fig. 1.**
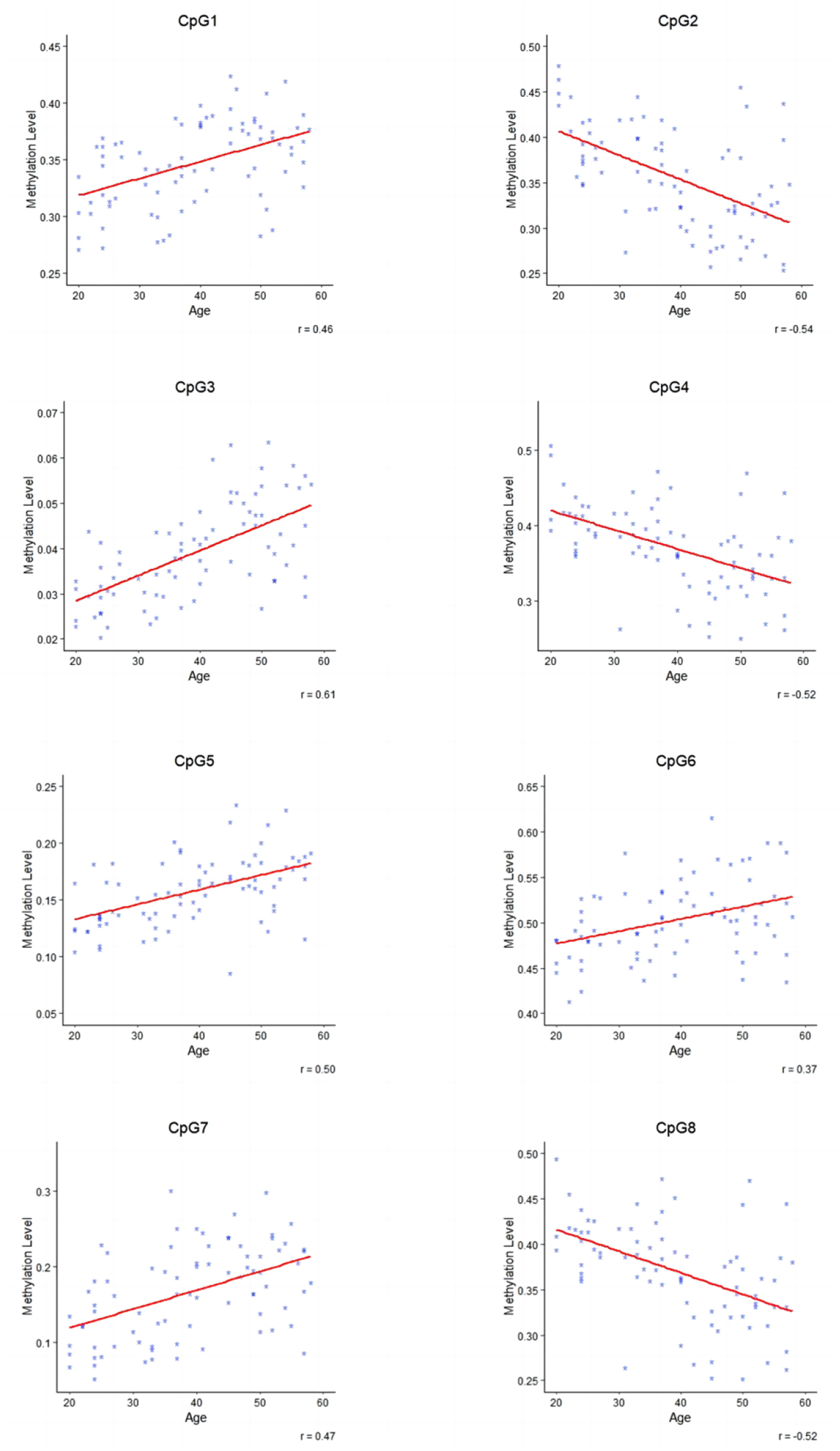
Regression scatter plot of DNA methylation levels at 5 CpG sites in blood samples and sample age.

**Tab.3.** R-value corresponding to candidate sites.

| Site | R |
| --- | --- |
| CPG1 | 0.46 |
| CPG2 | -0.54 |
| CPG3 | 0.61 |
| CPG4 | -0.52 |
| CPG5 | 0.50 |
| **CPG6** | 0.37 |
| **CPG7** | 0.47 |
| **CPG8** | -0.52 |

### 3.2 Site Selection and Data Processing

Based on Pearson correlation analysis, CpG2, CpG3, CpG4, CpG5, and CpG8 showed |r| values above 0.5, indicating a strong positive correlation between methylation levels and sample age. However, CpG8’s numerous outliers led to its exclusion from the study. Outlier samples were identified and excluded from the training set, ensuring robust model performance. Details are provided in the Tab.4.

**Tab.4.** Abandoned blood sample information.

| Sample number | Age | Age stage |
| --- | --- | --- |
| DZ-484 | 31 | 20~29 |
| JXH061 | 42 | 40~49 |
| JXH080 | 45 | 40~49 |
| JXH138 | 45 | 40~49 |
| JXS001 | 46 | 40~49 |
| L00341 | 50 | 40~49 |
| L00490 | 51 | 50~59 |
| L00398 | 57 | 50~59 |
| L00445 | 57 | 50~59 |
| L00445 | 57 | 50~59 |

### 3.3 Construction of Blood DNA Methylation Age prediction Model

Using data from 70 selected samples, a model was built based on CpG2, CpG3, CpG4, and CpG5 sites. A mathematical formula for age prediction was derived using SVR regression, considering methylation levels at these sites.

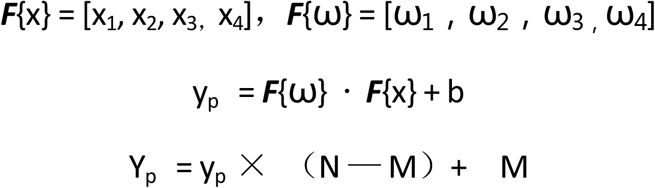

The model was trained and tested using MATLAB R2022a, yielding an MAE of 2.77 years and an accuracy of approximately 91.56% for the test set. The model’s performance was validated through cross-validation, ensuring its stability and accuracy. The selection of training model parameters and formula variable values is shown in Tab. 5.

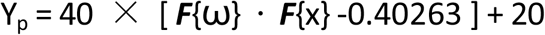

**Tab.5.**
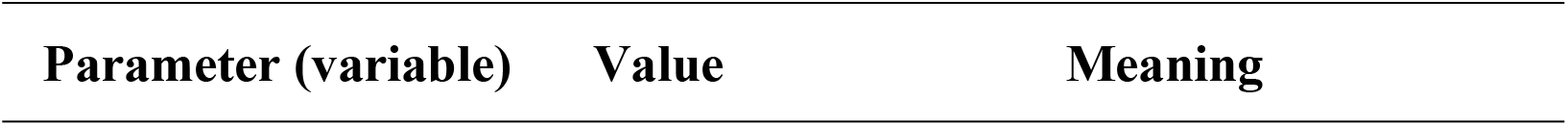
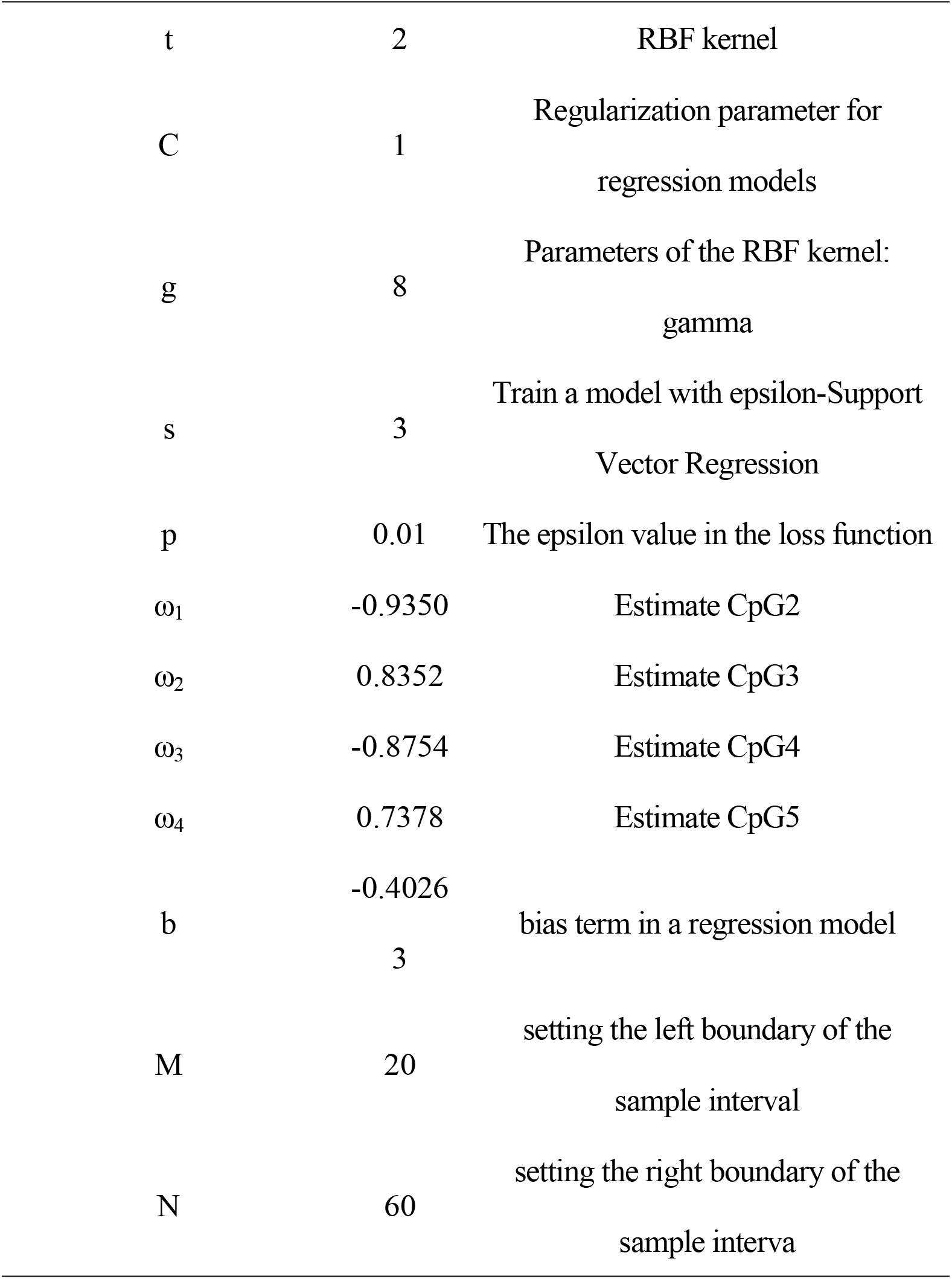
Training model parameters and formula variable values.

According to the calculation formula of relevant indicators, the training set R2 of the established SVR regression model is 0.90808, and the testing set R2 is 0.88529. Therefore, the age inference model constructed based on the CpG2-CpG3-CpG4-CpG5 locus combination can achieve an accuracy of about 89% for age prediction inference values, with a MAE value of 1.58 years for predicting age in the training set and actual age; The MAE value between the predicted age and the actual age in the test set is 2.57 years old. In the SVR regression model constructed this time, the fitting line graph between the predicted age of the training set and the true age is shown in Fig. 2, and the fitting line graph between the predicted age of the testing set and the true age is shown in Fig. 3.

**Fig. 2.**
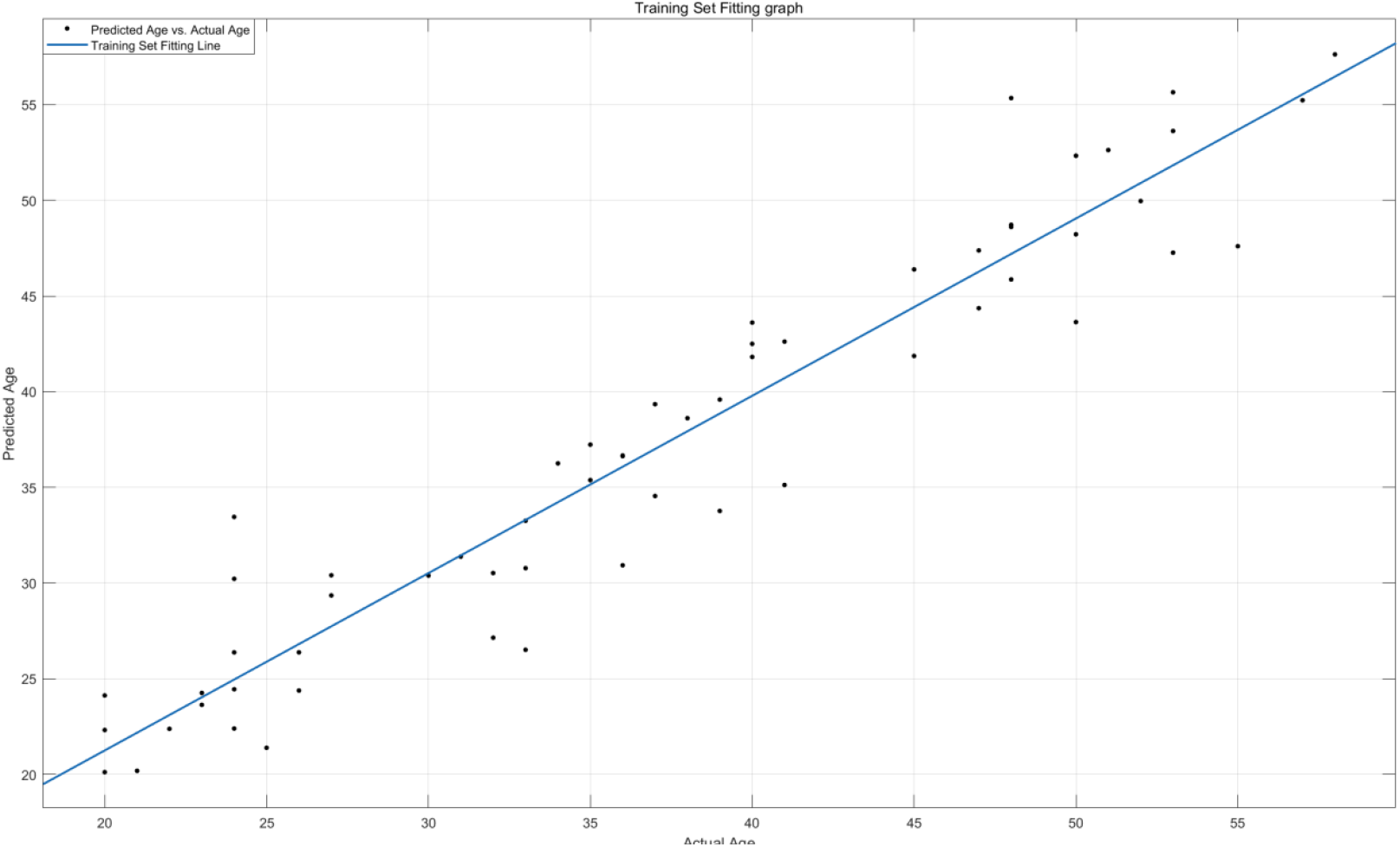
Fitting line graph between predicted age and actual age in the training set.

**Fig. 3.**
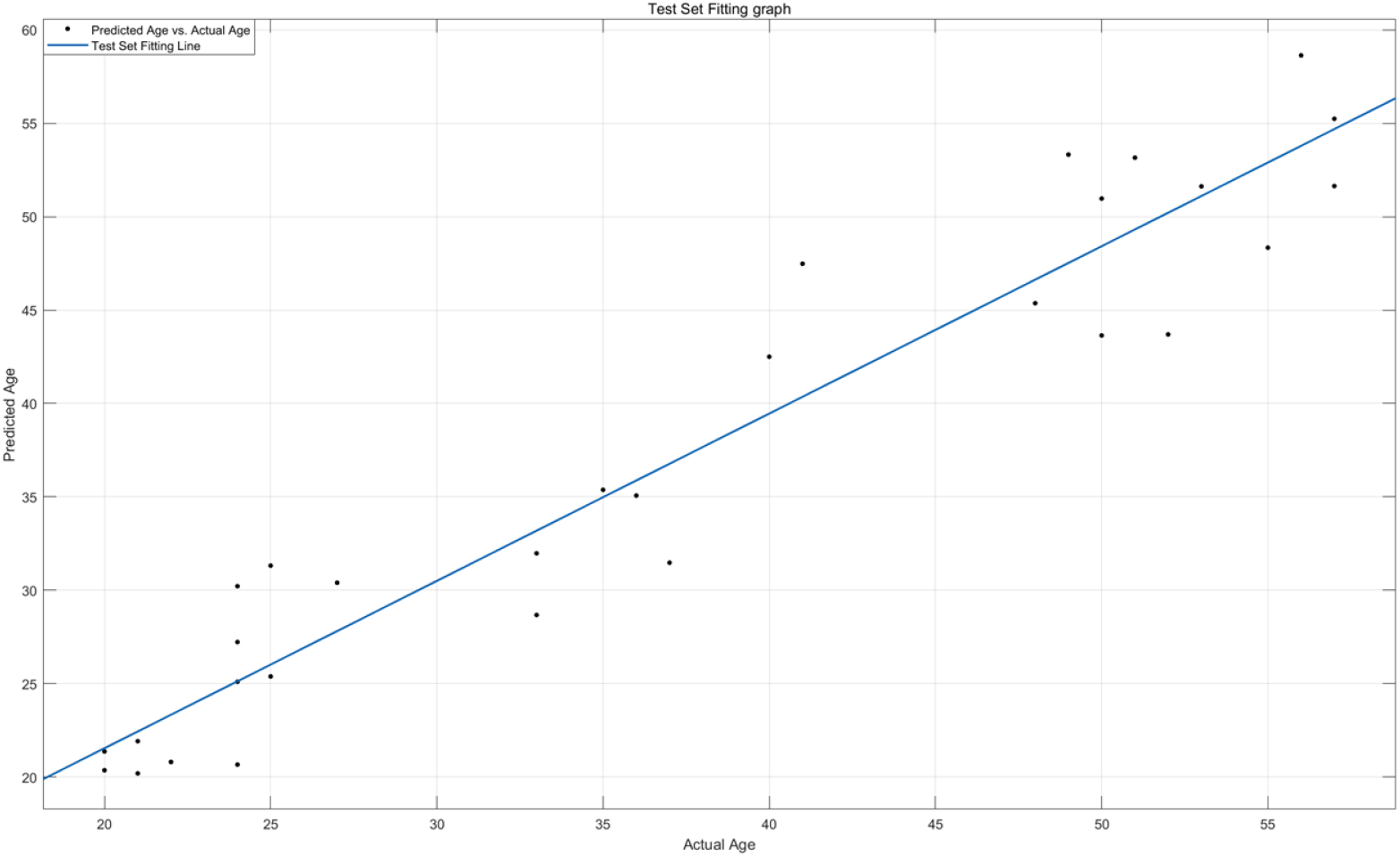
Fitting line graph of predicted age and actual age in the test set.

### 3.4 Accuracy Analysis across Different Age Groups

The model’s accuracy was assessed across four age groups using 60 training samples and 31 test samples. Performance metric ‘Accuracy’ are presented in Tab. 6 and Tab. 7 respectively. Despite decreasing Accuracy values with age group, indicating reduced predictive capability for older ages, the model demonstrated robust accuracy within ±5 years.

**Tab.6:** Accuracy metrics for age groups in the training set.

| Age groups | Accuracy | Number of samples |
| --- | --- | --- |
| 20-29 | 94.1% | 17 |
| 30-39 | 88.9% | 18 |
| **40-49** | 84.6% | 13 |
| **50-59** | 83.3% | 12 |

**Tab.7:** Accuracy metrics for age groups in the test set.

| **Age groups** | **Accuracy** | **Number of samples** |
| --- | --- | --- |
| **20-29** | 100% | 9 |
| **30-39** | 100% | 8 |
| **40-49** | 75% | 7 |
| **50-59** | 57.1% | 7 |

## 4. Discussion

Recent studies on DNA methylation age inference have increasingly focused on using a small number of highly linear CpG sites for model construction. In this study, CpG2, CpG3, CpG4, and CpG5 showed optimal correlation (r-values) with sample age, facilitating SVR model construction.

Through iterative training and refinement, a robust SVR mathematical model was established, achieving a 89.28% accuracy rate with a MAE of 2.77 years for age prediction. The model exhibited high stability and avoided overfitting, maintaining a self-check accuracy rate of 90%. However, as noted, MAD values increased with older age groups, suggesting a decline in predictive accuracy for older individuals.

